# SANE: an Index of Anthropogenic Noise Levels for Wildlife Research in Terrestrial Ecosystems

**DOI:** 10.64898/2025.12.23.696194

**Authors:** Matteo Giuliani, Davide Mirante, Luca Francesco Russo, Andrea Zampetti, Luca Santini

## Abstract

1. Understanding the impact of noise pollution on wildlife is challenging due to the complexity of isolating the anthropogenic components from the soundscape. Soundscape studies usually employ acoustic indices that apply arbitrary thresholds to isolate anthrophony from natural sounds. However, natural and anthropogenic sounds do not always conform to these thresholds, hindering both accuracy and comparability of the resulting indices. While significant progress has been made in automated identification of acoustic events through artificial intelligence-based classifiers, an effective method to quantify anthropogenic acoustic pressure is still lacking.
2. We propose a novel index, the Selective Anthropogenic Noise Exposure (SANE), which leverages BirdNET deep neural network to isolate human-related sounds from recordings. The index consists in the sum of the Median Amplitude Index of all human noise categories, thereby capturing both the intensity and cumulative impact of multiple disturbance events, while also enabling the decomposition of noise level across different categories of disturbance (e.g., traffic, human voices).
3. We test SANE’s performance in a real urban setting and through soundscape simulations and compare it with two frequency-based indices and another artificial intelligence-based acoustic index. SANE was effective in describing human noises within areas characterized by different levels of urbanization, improving upon other indices’ shortcomings. Additionally, SANE was robust at very fine temporal scales, precisely quantifying anthrophony levels for single recordings. Among the indices considered, SANE was the only one that remained insensitive to low-frequency biophony, which confounded both frequency and artificial intelligence-based metrics.
4. By leveraging artificial intelligence-based classifier’s ability to detect multiple human-made sound classes, SANE index captures the intensity of anthropogenic noise while being insensitive to non-conforming natural sounds to the traditionally identified anthrophony range. Furthermore, SANE has the potential to assess the relative contribution of different noise types to the overall acoustic pollution, opening new research avenues on the acoustic pollution effects on wildlife in high disturbance contexts.

## 1. Introduction

An increasing number of studies have highlighted negative effects of anthropogenic noise (anthrophony) on wild fauna ranging from behavioural to physiological alteration (Brown *et al*., 2012; Fuller *et al*., 2007; Shannon *et al*., 2016; Slabbekoorn & Peet, 2003; Slabbekoorn & Ripmeester 2008; Rosa & Koper 2018). Effects include the disruption of predator-prey interactions (Fortin *et al*., 2004; Sweet *et al*., 2022;de Jong *et al*., 2020; Kaiser & Hammers, 2009), changes in communication efficiency (Hödl, 1977) with subsequent adjustments of frequency range (Lampe *et al*., 2014; Loss *et al*., 2021), and ultimately consequences for individual fitness (Cruz *et al*., 2018; Jong *et al*., 2017). With the projected increase in human population and related infrastructures, anthrophony levels are expected to further increase in the near future (Hildebrand, 2009), underscoring the need for robust tools to quantify and monitor such disturbances.

Acoustic indices have recently gained popularity for their capacity of describing the soundscape with relatively low effort (Alcocer *et al*., 2022). Among the numerous indices that currently exist, some have been specifically designed to quantify anthrophony, i.e., the Power Spectral Density (PSD) calculated between 1-2kHz, and to measure the interaction between human noises and biophony, i.e., the Normalized Spectral Dissimilarity index (NDSI; Kasten *et al*., 2012). These indices rely on theoretical frequency boundaries that delimitate biophony (2-11 kHz) and anthrophony (1-2 kHz) (Pijanowski *et al.,* 2011). However, the frequency boundaries of soundscapes components are not as rigid as formalized, with many sounds exceeding their theoretical limits (e.g., Jacewicz et al., 2023; Sánchez-Giraldo et al., 2020).

Anthropogenic sounds can be highly diverse and produce different effects on species depending on their frequency and intensity (Gomes *et al.,* 2022; Magnotti & Luther, 2014). Similarly, biophony encompasses a huge variety of sounds, some of which fall within the anthrophony range, e.g., those produced by the Eurasian wild pig or by the yellow-legged gull. Thus, the ambiguity and the complexity involved in distinguishing between natural and anthropogenic sound represent a significant challenge, limiting the effectiveness of acoustic indices based on fixed frequency threshold. In this context, advanced methods for automated sound analysis can be a valuable support. Artificial Intelligence (AI) is increasingly applied in ecology to support and accelerate data processing (Pollock et al., 2025). In the field of bioacoustics, several algorithms based on Neural Networks have been applied to identify species from acoustic recordings (e.g. BirdNET; Kahl *et al*., 2021), or to discriminate between anthrophony and biophony (e.g. CityNet; Fairbrass *et al.,* 2019).

Here we present a novel index, the Selective Anthropogenic Noise Exposure (SANE), which exploits AI to isolate and quantify the amplitude of anthropogenic noise from acoustic data to estimate anthropogenic noise levels for impact assessments and biodiversity research. We illustrate how SANE performs in describing the intensity and identity of anthropogenic noise in two case studies and compare it with the anthrophony activity level estimated by CityAnthroNet (Fairbrass *et al*., 2019), and with frequency-based indices, i.e., NDSI and PSD (1-2kHz). First, we took advantage of an extensive passive acoustic monitoring conducted in the city of Rome (Italy), spanning from large peripheric parks to the urban centre. We visualized the spatial and temporal pattern of all calculated indices to evaluate their consistency and overall performance in capturing anthrophony levels. Furthermore, we showcase SANE’s decomposability by using the values obtained for each human noise category, highlighting their relative temporal and spatial variation along the urbanization gradient. Second, we test indices’ abilities to describe anthrophony using simulated soundscapes mimicking gradients of noise pollution, and varying proportion of bird species vocalizing within the range of biophony and anthrophony.

## 2. Methods

### 2.1. BirdNET as a classifier of human noise

BirdNET is a Deep Neural Network originally designed for automated birds’ species identification, which is widely used in ecological studies that employ acoustical data (e.g., Plat *et al.,* 2025; Paterson *et al*., 2024; etc.). In addition to ∼1000 bird species, BirdNET is also capable of recognizing nine classes of anthropogenic sounds; specifically, human vocal, human non-vocal, a class encompassing several sounds directly produced by humans (sneezes or stepping, human whistle, engine, power tools, fireworks, siren, guns) and dogs (Kahl *et al.,* 2012). The human noise training dataset consisted of almost 8000 recordings derived from the Google AudioSet (Gemmeke *et al*., 2017), the Freefield1010 (Stowell and Plumbley, 2013) and the WarblR datasets, encompassing audio of worldwide provenance (Kahl *et al*. 2021).

Other deep learning systems have been specifically designed to quantify anthropogenic noise, like CityAnthroNet (Fairbrass *et al*., 2019). However, CityAnthroNet training dataset is geographically restricted, with recordings collected exclusively across the Greater London area (England, United Kingdom). Furthermore, it does not discriminate between disturbance classes, and its default output returns the proportion of human noise activity throughout the entire recording, without estimating the intensity of anthrophonic sounds.

### 2.2. SANE index calculation

The Selective Anthropogenic Noise Exposure index (SANE) is calculated in 3 steps (Fig.1). Firstly, acoustic files are processed by BirdNET to identify human related sounds. BirdNET divides the audio file in snippets of 3 seconds and classifies each in a sound category. So, all snippets classified as human related sounds are identified. BirdNET also provides a confidence score associated with each sound. Then, the optimal confidence threshold to obtain the desired precision should be set based on a local validation (see section 2.4.2). Secondly, snippets classified as human related sounds are cut and used to calculate the Median Amplitude index (Depraetere *et al*., 2012), i.e., the median of the amplitude envelope, which is not computational demanding and quantifies the amount of acoustic energy associated with each anthropogenic acoustic signal identified without being sensitive to outliers. Thirdly, Median Amplitude Index values estimated from each snippet are summed for each audio to produce a SANE value for each human noise category, as well as a global SANE value considering all human noise categories. If no human noise was recorded throughout the entire recording, the SANE index equals to zero. The described workflow is embedded in a R customizable function, which is provided through a GitHub repository ([link omitted for anonymous peer review, consult the link provided in the Data Accessibility Statement to temporarily access functions and data for peer review only]).

**Figure 1.**
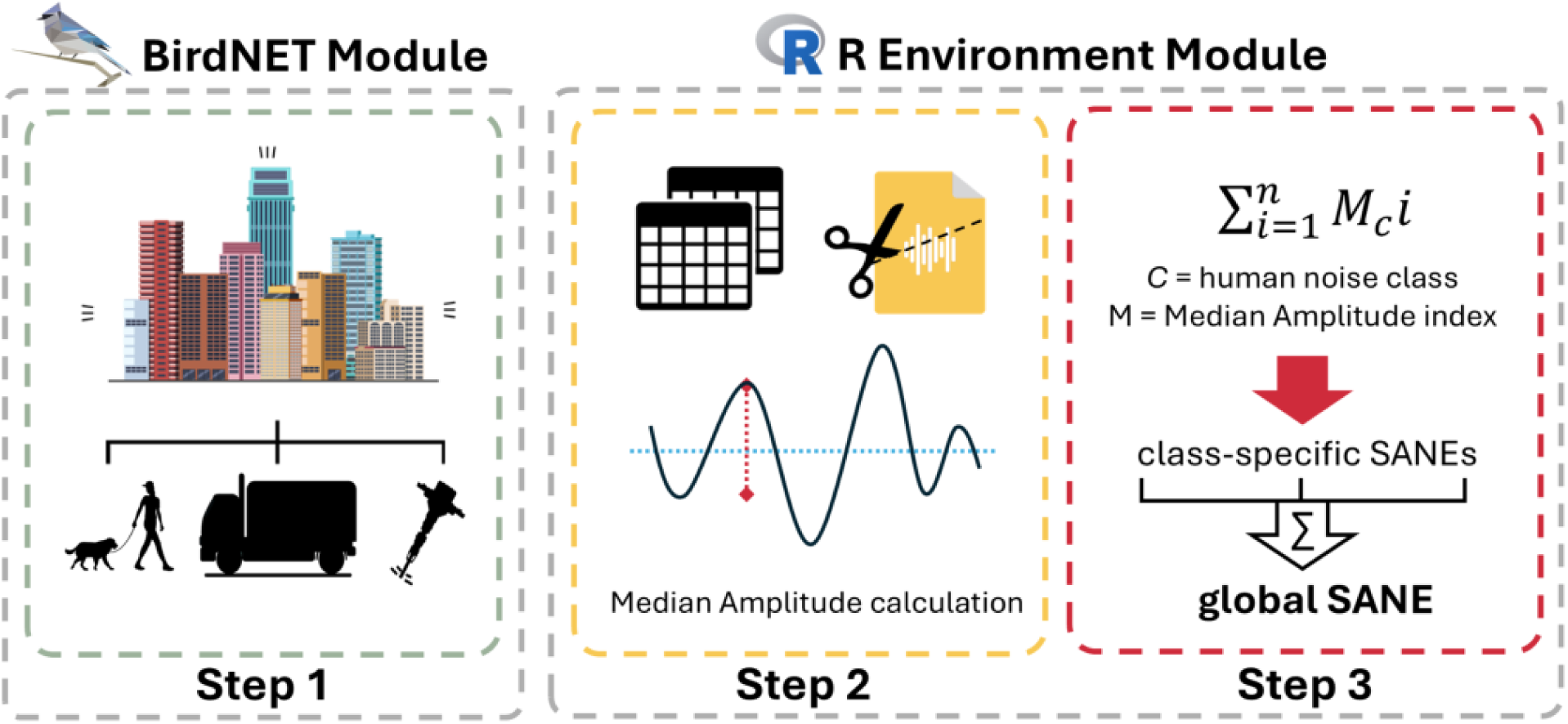
Workflow of the SANE index calculation. The first step uses BirdNET to identify target human noise signals. Thus, index processing moves in R: specifically, in step 2 the BirdNET output is used to compute Median Amplitude for each signal; in Step 3, Median Amplitude values are summed within and across noise classes to obtain the SANE indices of interest.

To assess the robustness of the SANE index relative to alternative approaches and illustrate its application, we apply it both to real-world recordings (2.3), and to controlled simulated soundscapes (2.4).

### 2.3. Real soundscape tests

#### 2.3.1. Acoustic survey

We took advantage of an extensive passive acoustic monitoring taking place from March to June 2024 in the city of Rome (Italy). Rome is among both the largest and greenest cities in Europe, with 49% of its large area covered by green spaces (Russo et al., 2025). This diverse landscape offers a highly heterogeneous soundscape and an excellent opportunity to study gradients of anthropogenic sounds. The survey consisted of a total of 75 listening points within the motorway (Grande Raccordo Anulare) encircling the most populated and urbanized part of the city (Fig. S1). The minimum distance between each listening point was no less than 100 m to avoid spatial pseudoreplication (Alcocer et al. 2022). To characterize the urban environment, we used the map provided by Russo et al. (2025) that classifies the landscape into three urbanization categories: Low Urbanization representing homogenous green spaces, Moderate Urbanization where impervious surfaces are interspersed with small green patches, and High Urbanization representing highly impervious areas.

Each listening point consisted of one AudioMoth v1.2.0, which remained in place one night per month from 5:00 PM to 9:00 AM recording for 5 minutes every 20 minutes (5 hours and 20 minutes of recording per day). The deploying period was designed to intercept the vocal activity peaks of diurnal and nocturnal birds, while minimizing the risk of device theft. The final set consisted of a total of 13,100 five-minutes recordings, 64 per day of deployment for each listening point, for a cumulative recording time of 65,500 minutes.

#### 2.3.2. BirdNET validation

We used BirdNET to identify the human noise signals in each audio file of all recordings collected (∼65,000 min.). We compiled an *ad hoc* species list comprising solely the full set of human activity-related sounds encompassed by BirdNET for audio-analysis. BirdNET processes audio files in 3-second snippets, assigning a confidence score when a target sound is identified.

We set the minimum confidence score threshold at 0.1, applied a 2s of overlap on prediction segments to enhance recall (Pérez-Granados *et al*., 2025), and maintained the default sensitivity (1.0).

To ensure that only target-sounds were classified and to avoid errors from false positives, which would affect index’s accuracy, we validated the outputs. We followed the BirdNET validation procedure proposed for birds’ sounds by Pérez-Granados (2023); thus, we randomly selected 100 classified audio segments from the BirdNET output for all the classes of anthropogenic noise, ensuring an even distribution across all confidence score intervals (e.g., 0–0.2, 0.2–0.4,…, 0.8-1). Each segment was manually reviewed and annotated to confirm the presence of human-related noise. Annotations indicated whether the segment truly contained anthropogenic sounds (true positive) or not (false positive), as well as whether the detected sound matched the class assigned. We then computed the precision score, defined as the ratio between true positives and the sum of true and false positives. This was done for each threshold considered (i.e., thresholds ranging from 0 to 1 in steps of 0.05), both for each anthropogenic noise class separately and anthrophony overall. Furthermore, we also assessed the recall. Specifically, we randomly selected five recordings that contained at least one recognized signal of a specific class, for a total of 45 5-minutes recordings. Every recording was listened to, and human sounds were manually annotated for each 3-seconds snippet. Then, we calculated the recall as the ratio between true positives and the sum of true positives and false negatives. Lastly, we also computed the F1 scores as a balanced measure of precision and recall, defined as twice the ratio between the product and their sum.

#### 2.3.3. Anthrophony estimates through SANE and other indices

To calculate SANE indices on our acoustic dataset, we followed the three-step procedure described above (Figure 1). We used the precision metrics of the classifiers to identify the optimal thresholds to avoid false positives. We used precision instead of F1 score for threshold selection to reduce the chance of including non-target sounds in our index, thereby overcoming a common problem of frequency-based indices. We considered only the human audible frequency range (0-20kHz). We then obtained one SANE value per sound class and one global SANE value per recording.

For each recording, we also calculated the CityAnthroNet level of anthropogenic acoustic activity (Fairbrass *et al.,* 2019), the Normalized Difference Soundscape Index (NDSI), and the Power Spectral Density (1-2 kHz, PSD) as described by Kasten et al. (2012). CityAnthroNet classifier was run in Python v.2.7.12 (Python Software Foundation, 2016) using scikit-learn v.0.18.1 (Pedregosa et al., 2011) and matplotlib v.1.5.1 (Hunter, 2007), as designed by Fairbrass *et al*. (2019). The NDSI was defined as the ratio ([biophony-anthrophony]/[biophony + anthrophony]) of the estimated PSDs, computed using Welch’s method (Welch, 1967), of the anthrophony (1–2 kHz) and biophony (2–11 kHz). Acoustic indices computation procedures were conducted in R studio (R Core Team, 2024) using the *seewave* package (Sueur et al., 2008a, Sueur et al., 2008b).

### 2.4. Simulated soundscapes tests

To evaluate the ability of acoustic indices in capturing anthrophony while controlling for confounding variables, we produced 100 simple simulated soundscapes under 4 scenarios, containing only human noises and birds’ vocalization. We designed the simulated soundscapes to be temporally structured as a sequence of contiguous, non-overlapping 3- seconds audio segments, with no silent gaps. We retrieved the sound snippets from a small classified acoustic dataset, consisting in ornithological recordings, collected from xeno-canto https://xeno-canto.org), and anthropogenic sounds with low background noise, extracted from the field case study (section 2.3; see Table S1 for details). We classified audio segments in our dataset into three classes: non-anthrophonic birds, i.e., avian vocalizations that do not fall into the anthrophony frequency range (1-2 kHz), anthrophonic birds, i.e., birds’ signals that overlap with the anthrophony frequency range, and human noise.

Each simulated soundscape had a total length of 1200 seconds, divided into four 300-second phases, each with a distinct snippet class selection: (*i*) only *non-anthrophonic birds* and *human noise*; (*ii*) only *anthrophonic birds* and *human noise;* (*iii*) both *anthrophonic* and *non-anthrophonic birds*, and *human noise*; (*iv*) only *anthrophonic* and *non-anthrophonic birds*. In the first three phases (*i-iii*), the proportion of bins filled with *human noise* increases by 1/10 every 30 seconds starting from zero. In the last phase (*iv*) anthrophonic bird snippets increased progressively, following the previous human noise pattern.

Finally, to generate soundscapes that were not influenced by the source recordings’ audio signal intensity, we normalized each snippet to a fixed amplitude (1.0). We produced the simulated soundscapes in R (R Core Team, 2024), using the *tuneR* (Ligges *et al*., 2023) and the *seewave* (Sueur et al., 2008a, Sueur et al., 2008b) libraries.

As for the real soundscapes, we computed the global SANE indices, using the precision metrics to guide the choice of confidence threshold to achieve the maximum precision possible (1), the CityAnthroNet level of anthropogenic acoustic activity, the NDSI and the PSD (1-2 kHz). We did not calculate the indices for each entire recording; instead, we computed them for each 30-seconds segment to reflect the sequential structure of the soundscape and to ensure a fine-scale assessment of index responses to increasing snippet frequencies of human noise (phases *i-iii*) and anthrophonic bird (phase *iv*).

### 2.5. Indices comparison

For real soundscape recordings, we computed 8 indices: five SANE indices (a global one and one for each human noise category retained), CityAnthroNet activity level, PSD (1-2 kHz) and NDSI. This way our dataset provided a detailed temporal series of the values assumed by the indices computed through our sampling period in each site. For temporal series estimation, we grouped sites on the basis of their Urbanization category (see 2.1). For direct comparability among indices, we scaled them to a mean of zero and standard deviation of 1. Furthermore, because higher values of the NDSI indicate lower anthropogenic presence, for enhancing interpretability we used the negative form of the NDSI (invNDSI).

For simulated soundscapes, we aimed to evaluate the consistency of each acoustic index in capturing anthropogenic noise under controlled conditions. We summarized the behaviour of each index across simulated soundscapes by calculating the mean value for each 30-second time step. We then produced a time series plot of the indices using the scaled mean values (mean = 0, SD = 1) of all indices for direct comparison and evaluated the indices variability across simulations.

Our dual approach, combining real and simulated soundscapes, enabled a comprehensive evaluation of index performance. Specifically, it allowed us to assess each indices’ ability to capture temporal dynamics of anthropogenic noise in complex environments, to discriminate among urbanization gradients, and to maintain consistency across repeated, controlled soundscapes.

## 3. Results

### 3.1. BirdNET classifier’s performance

Within the complete audio material, BirdNET identified a total of 434,865 human noises, i.e., 3-seconds segments containing an acoustic signal of human activities. For BirdNET, the most represented human noise class was “Engine”, which amounted to 73% of the total.

Conversely, the less frequent class was “Human Whistle”, consisting only of 28 detections. A relatively frequent disturbance class, “Gun”, consisted only of false positives.

BirdNET’s output validation process highlighted a notable ability to discriminate between natural (e.g., birds’ songs) and anthropogenic noise: the mean precision value throughout the confidence thresholds was 0.99 for BirdNET, with a minimum value of 0.97 for the 0.1 threshold. The minimum confidence threshold that maximized precision was 0.5 (Fig. 2).

**Figure 2.**
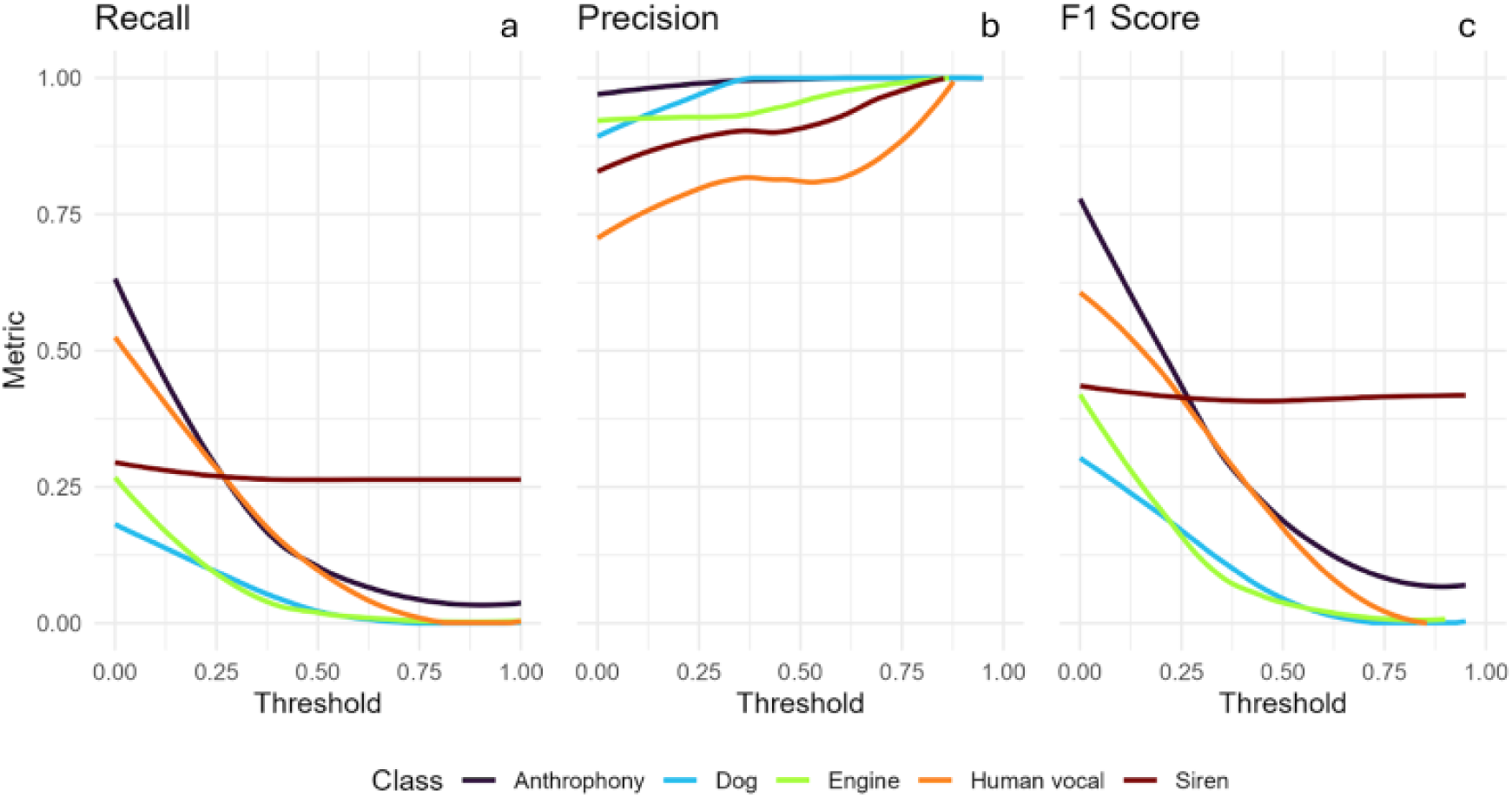
Ranges of recall, precision, and F1-score for selected human noise classes identified by BirdNET across threshold values. Metrics were computed for the four most represented classes (i.e., Dog, Engine, Human Vocal, and Siren) as well as for the global Anthrophony category.

The performance of BirdNET in classifying human noise as anthrophony varied across classes (Fig. 2). Several classes resulted to be quite rare, making the computation of their metrics challenging; thus, we considered them for the global anthrophony, but we did not compute neither their precision, recall and F1-score. Classes that were sufficiently represented were Human Vocal, Dog Barking, Engine and Siren.

Recall values varied for all disturbance classes, with the general anthrophony exceeding 0.6 at the 0.1 confidence threshold (Fig. 2a). Due to the high precision, the classifier obtained a relatively high F1-score for the general Anthrophony at the minimum Confidence threshold possible (Fig. 3c).

**Figure 3.**
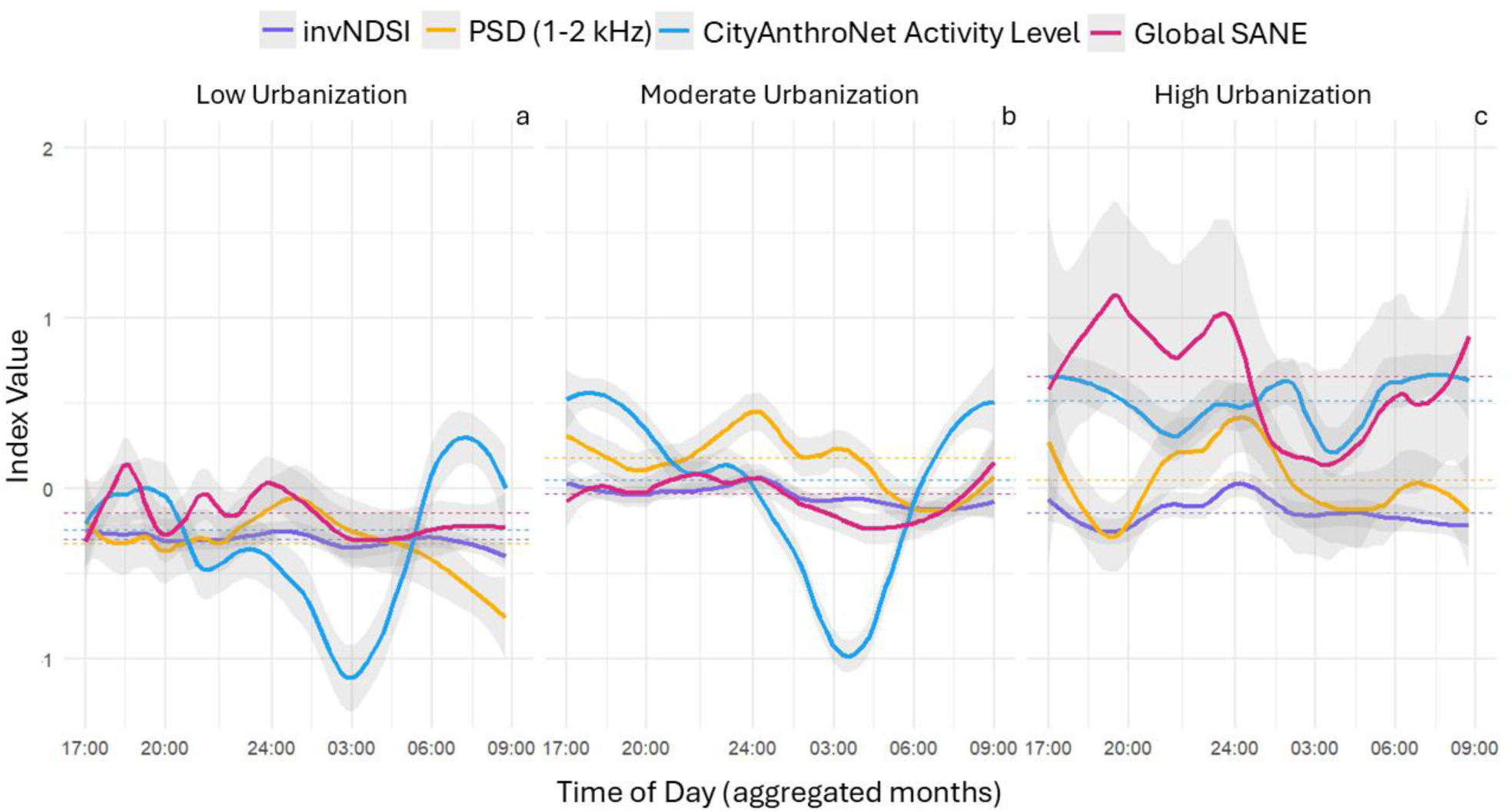
Time-series plot of the invNDSI, the PSD (1-2 kHz), the CityAnthroNet Activity Level and the global SANE through urbanization cluster obtained by aggregating together the data from all the sampling months and using the *Loess Regression* approach (*span* = 0.6). The shading around the curves represents the 95% confidence intervals. All indices were scaled to mean 0 and standard deviation of 1. Dotted horizontal lines represent the mean values of each index in each urbanization class.

### 3.2. Real Soundscape temporal patterns of anthrophony

Frequency-based indices showed similar temporal patterns within different levels of urbanization, with PSD displaying larger fluctuation. On the contrary, AI-based indices were markedly distinct.

The temporal trends in the four indices revealed interesting patterns in relation to the increase in human activity and biophony around sunset and sunrise hours, across the three urbanization levels (Low urbanized = green areas, Morderate urban = impervious areas interspersed with green areas, High urban = impervious areas; Fig. 3). The SANE was the only index highlighting an increase in intensity in high urbanized areas during evening hours, reflecting late afternoon traffic and night life. Instead, the CityAnthroNet showed a higher activity in low and moderate urban areas than in high urbanized areas during late hours, probably reflecting the sunset peak in biophony. Frequency-based indices did not show consistent increases across the three urbanization levels in the early morning. On the contrary, both AI-based indices tended to increase in the morning, except for the SANE who does not in low urbanized areas. Interestingly, the CityAnthroNet activity level increases substantially in the morning also in the low urbanized areas.

When the SANE index was decomposed into different noise classes (Fig. 4), it revealed that low-urbanized areas were dominated by human voices; low and moderately urbanized areas by traffic noise and dog barking; and moderately to highly urbanized areas by sirens.

**Figure 4.**
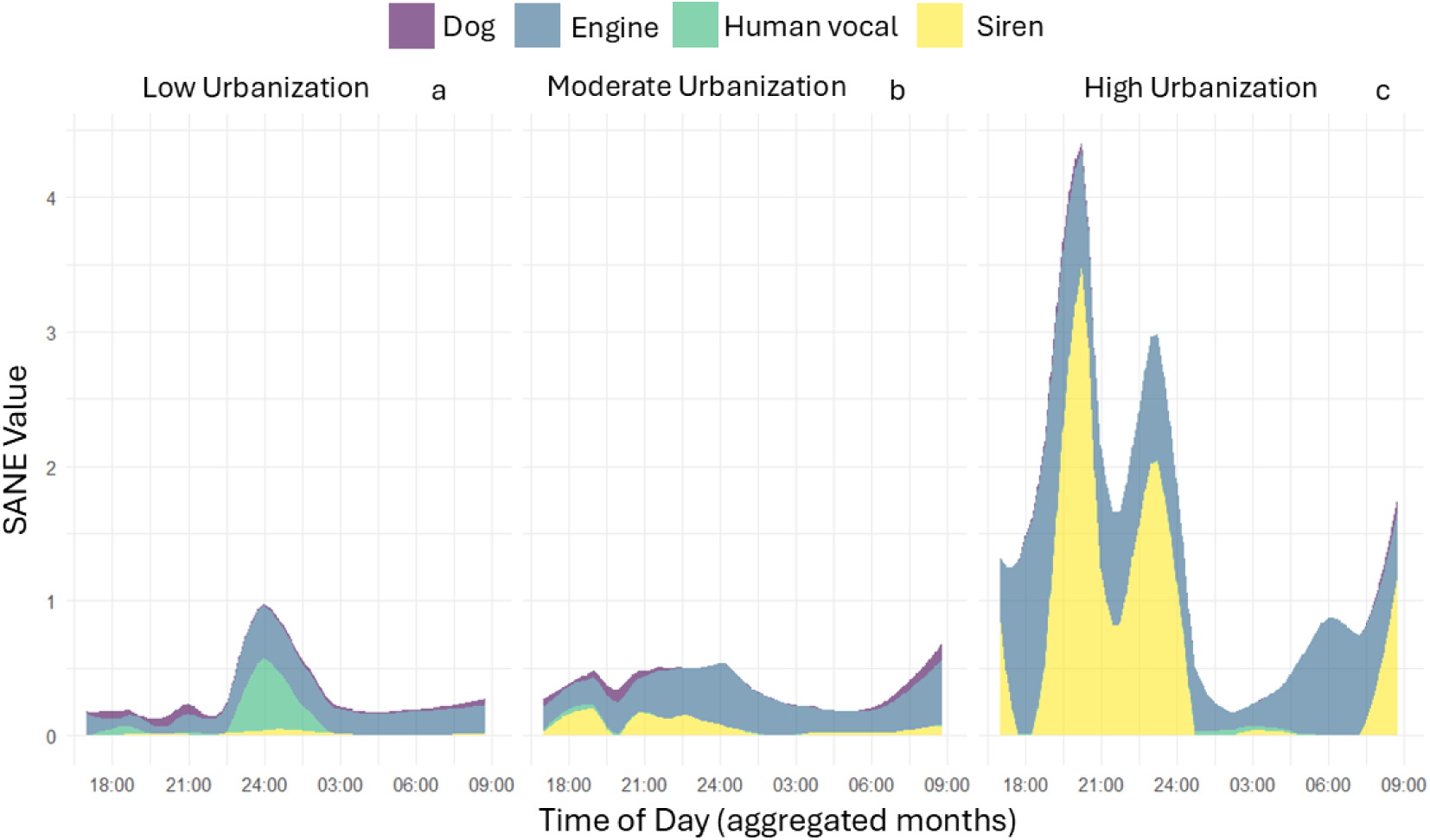
Area chart representing the contribution of each human noise class to the global SANE through the time series, obtained aggregating months, within different urbanization levels. To enhance the visualization of temporal trends of SANE values, we applied a locally weighted smoothing technique by using the *Loess* approach. The *Loess* was performed only for groups with at least ten detections with a smoothing span of 0.6, being coherent with the time-series plot. Thus, three disturbance classes are missing, specifically, the “Gun” class, due to the absence of true positives, the “Human-whistle” class, for having too few detections, and the “Human non-vocal” for the impossibility of achieving a 90% precision. The upper line of each area represents the value assumed by the global SANE.

### 3.3. Simulated Soundscape Acoustic Indices patterns and consistency

The SANE index was the least affected by the presence of anthrophonic bird vocalization across the 100 simulated soundscapes. During the first three scenarios (Fig. 5*i*-*iii*), SANE values increased proportionally to the number of bins occupied by human noises in each 30 second step; in the last scenario, which only included bird sounds (Fig. 5*iv*), its pattern remained low and flat. In contrast, frequency-based indices were generally higher in the second scenario including anthrophonic bird sounds (Fig. 5*ii*) than in scenarios one and three (Fig. 5*i*, *iii*), resulting in a less marked increase relative to the actual rise in anthropogenic noise snippets frequency. In the fourth scenario, both indices increased in response to the growing presence of anthrophonic birds.

**Figure 5.**
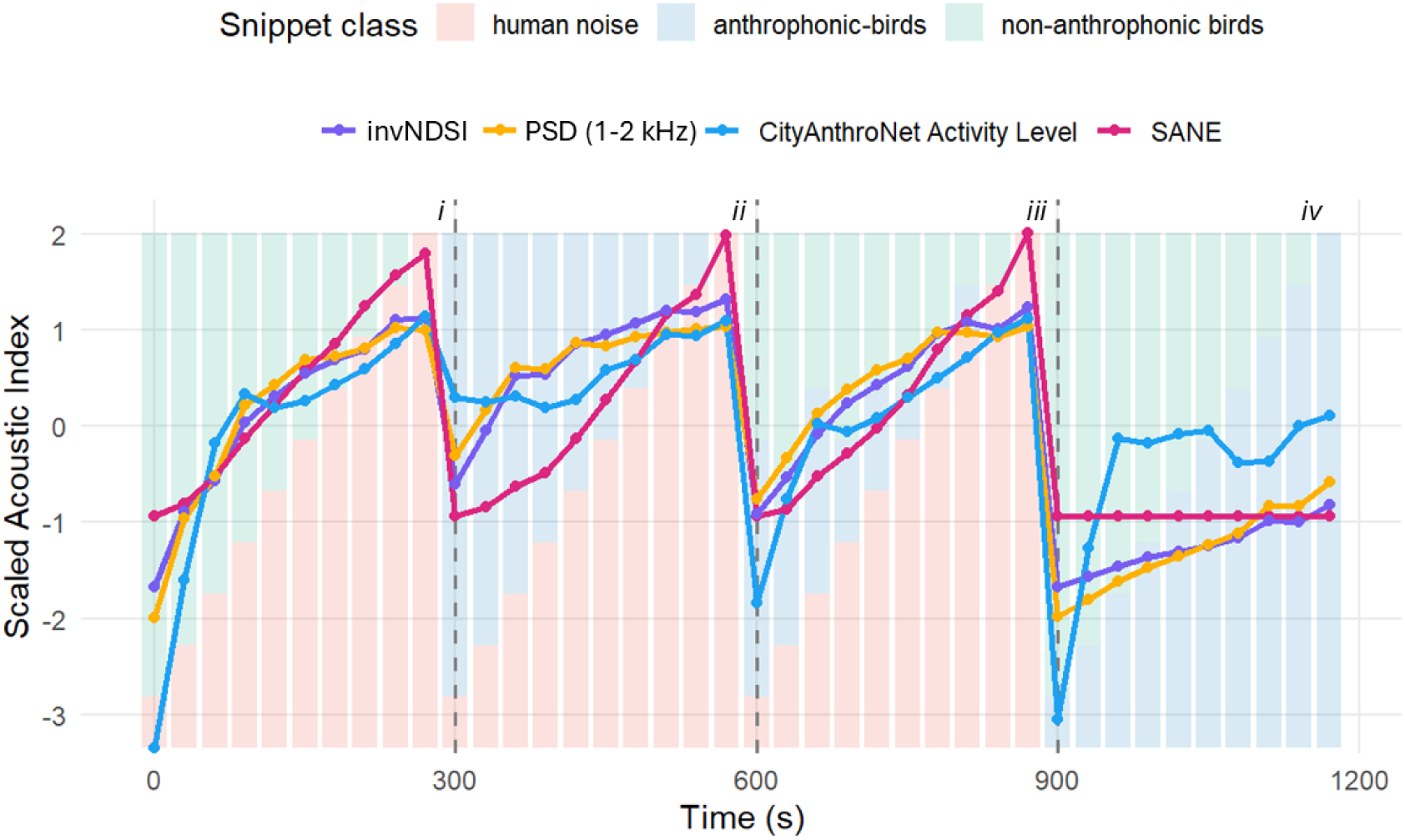
Temporal patterns of the invNDSI, PSD (1–2 kHz), CityAnthroNet Activity Level, and global SANE, averaged across the 100 simulated soundscapes. All index values were scaled to a mean of 0 and a standard deviation of 1. Points represent the mean value of each index for a 30-second time step; lines connect consecutive time steps to show temporal trends. Background bars represent the proportion of 3-second snippets constituting each 30-second time step appertaining to a specific class. Simulated soundscapes consisted in four sequential phases delimited in figure by the dashed lines: (*i*) only non-anthrophonic birds and human noise; (*ii*) only anthrophonic birds and human noise; (*iii*) comprehensive of anthrophonic and non-anthrophonic birds, and human noise; (*iv*) only anthrophonic and non-anthrophonic birds.

Surprisingly, the CityAnthroNet anthrophony activity level showed a pattern similar to the frequency-based acoustic indices, being higher in the second scenario including anthrophonic birds (Fig. 5*ii*) and increasing progressively in the fourth scenario lacking anthropogenic noise (Fig. 5*iv*). In particular, the increase in the last phase was sharper than those of the invNDSI and the PSD (1-2 kHz). Finally, AI-based indices displayed high consistency across the 100 simulated soundscapes (Fig. S2*a*,*d*), whereas frequency-based indices exhibited high variability under all scenarios (Fig. S2b,c).

## 4. Discussion

Here we introduced the Selective Anthropogenic Noise Exposure index (SANE), which combines an AI-based classifier with an acoustic index to quantify anthropogenic noise in terrestrial ecosystems. Deep learning tools make it possible to overcome one of the most important acoustic indices limits, i.e., the inability of isolating target sounds (Alcocer *et al.,* 2022). The application of an *ad-hoc* designed classifier to measure anthropogenic noise is not new in ecological studies (Fairbrass et al. 2019). However, the cumulative nature of the SANE index, together with its integration of median amplitude, provides a finer description of noise levels and outperforms indices based on the proportion of time in which anthropogenic sounds are detected (e.g. CityAnthroNet activity level).

### 4.1. SANE performance in real and simulated soundscapes

We found that frequency-based indices produced a counterintuitive result along the urbanization gradient, with both invNDSI and PSD (1-2kHz) being higher in Moderate than in High Urbanization sites. These inconsistencies likely stem from the inability of the frequency-based indices to avoid the influence of non-target sounds, introducing natural noise into the estimation of anthropogenic sound levels. Biological sounds falling within the anthrophony frequency range, i.e., 1-2kHz (Pijanowski *et al*., 2011), for example calls emitted by the Eurasian wild boar or by the European badger, can make undisturbed areas be recognized as acoustically polluted. However, not only biological sound can fall within the anthrophony range, several sources of traffic noise, such as car horns, high-revving engines, that contribute significantly to the overall human acoustic contribution to the soundscape, often exceed in the standard biophonic frequency range.

The lower invNDSI and PSD values observed in the early morning (Fig. 3a) can be explained by the presence of nocturnal non-target sounds. By inspecting the spectrogram, we noticed bat calls emitted in proximity of the recorder filling the entire spectrogram, hence falling into the anthrophony resulting in an inflation of NDSI and PSD. In contrast, the CityAnthroNet activity increased sharply in the morning followed by a decline in intensity, across the three urbanization classes. Such temporal pattern reflects the acoustic activity of bird choruses, which are known to be capable of dominating the soundscape (Ross *et al*., 2018; Staicer *et al*., 2019), hence being identified as anthropogenic sounds. Differently to the other indices, the global SANE consistently showed a progressive increase starting from the dawn across all urbanization classes. A further important difference between the SANE and the CityAnthroNet activity level can be clearly noticed from their pattern in the High Urbanization cluster (Fig. 3c). In the morning hours both indices tend to rise progressively, however only the CityAnthroNet activity level reached a plateau. This is ascribable to its mathematical formulation: when every second of a recording contains human noises that are recognized with high confidence, CityAnthroNet saturates and stabilizes to 1. Conversely, because the SANE index accounts for sound intensity it does not reach an upper limit.

All indices but the SANE were confused by the presence of anthrophonic birds. Minimum values of frequency-based indices were generally higher in the presence of anthrophonic bird sounds (Fig. 5), and both invNDSI and PSD (1-2kHz) and CityAnthroNet activity level increased with the increase of anthrophonic birds (Fig. 5*ii-iv*). While this was expected for frequency-based indices, it was not for CityAnthroNet. Yet, this might be explained considering this classifier’s training dataset, in which anthrophony represented more than 80% of sounds (Fairbrass *et al*. 2019). An imbalance in the training class sample size can produce a classifier biased toward the target class that flags inputs that only vaguely resemble the training data. Double-class classifiers, such as CityAnthroNet, are particularly sensitive to this issue (Wei & Dunbrack, 2013; Zheng & Jin, 2020). The SANE index was unique in remaining flat in the fourth time block of the simulated soundscapes (Fig. 5*iv*), recognizing the complete absence of target sounds.

SANE was also the most consistent across all simulated soundscapes in describing the level of human noise. Frequency-based indices, such as NDSI and 1–2 kHz PSD, necessitate hundreds of hours of recording before their values converge on a representative measure of the soundscape (Bradfer-Lawrence *et al*., 2019). As a result, their values on a single audio recording are often unreliable, as demonstrated by the high variability across simulated soundscapes (Fig. S2*a*&*b*). In contrast, the SANE was able to quantify disturbance at a very fine temporal scale, suggesting reliable application even at the single-recording level.

### 4.2. SANE decomposability

Another substantial advantage of the proposed approach is the possibility to characterize anthropogenic soundscapes by isolating different classes of anthropogenic sounds. Considering that different sources of noise can prompt responses of different nature and intensity in wild fauna (Francis *et al*., 2013; Shannon *et al*., 2016), the decomposability of the SANE index offers a considerable advantage over other indices. Depending on the focus of their investigations, researchers may be interested in isolating only a specific facet of the anthrophony or assess the diverse impacts of different human-related noises. For instance, Randler *et al*. (2006) has shown that dog barks are perceived as an immediate predatory risk by the Eurasian Coot, while McClure *et al*. (2013) has shown that motor vehicles are perceived as background noise by several bird species, causing acoustic signals masking. Furthermore, thanks to the decomposability, it is possible to determine which are the main components of the global SANE, and therefore which disturbance classes contribute the most to the level of human noise (Fig. 4). In our case study, for instance, we were able to determine that the main contributors to the global SANE values varied depending on the level of urbanization, with the Engine class being preeminent in both low and moderate urbanized areas, and the Siren class contributing the most in highly urbanized sites.

### 4.2. Potential Limits

For the human-related noise classes for which we were able to compute performance metrics, recall scores were relatively low, but precision remained consistently high. This is particularly important in describing ecological patterns, where false positives are often more undesirable than false negatives (Clare et al., 2020; Cowans et al., 2024; Guillera-Arroita et al., 2017). Thus, although some signals were inevitably missed, this loss did not distort soundscape patterns, because the high precision allowed us to reliably capture the temporal dynamics of the four retained classes.

## 5. Conclusion

The synergy between artificial intelligence and acoustic indices, here presented through the novel Selective Anthropogenic Noise Exposure (SANE) index, has shown to improve the quality of anthropogenic noise quantification compared to CityAnthroNet, and to frequency-based indices (NDSI and 1-2 kHz PSD).

By leveraging AI-based classifier’s ability to detect multiple human-made sound classes, SANE captures the intensity of anthropogenic noise, and it has the potential to assess the relative contributions of different noise types to the overall acoustic pollution. Furthermore, not being tied to a specific frequency range, SANE can also be computed only in a specific spectral window, like the human-audible range or the ultrasonic one. This SANE’s characteristic could open new research avenues, for example the study of the effect of ultrasonic anthropogenic noise on bats that remains unexplored (Li *et al*., 2025).

Here we have applied the SANE index on the BirdNET classifier, but the SANE workflow can be easily adapted to different classifiers. With the release of updated versions of BirdNET or new classification algorithms, its performance can further improve in the future. Thus, irrespective of the classifier used, this work emphasizes the importance of integrating artificial intelligence tools with acoustic indices to better assess the intensity and the nature of human noises, offering a reliable and flexible solution for acoustical research.

## Data availability

The function to calculate the SANE index is freely accessible on the GitHub repository at https://github.com/matpagle/sane. Moreover, users can directly import the function in the environment by calling s*ource(*https://github.com/matpagle/sane/blob/main/sanefunction.R) within R.

## Author Contributions

*Matteo Giuliani, Luca Francesco Russo, Davide Mirante, Andrea Zampetti and Luca Santini conceived the ideas and designed methodology; Matteo Giuliani collected the data; Matteo Giuliani, Luca Francesco Russo and Davide Mirante analysed the data; Matteo Giuliani and Luca Santini led the writing of the manuscript. All authors contributed critically to the drafts and gave final approval for publication*.

## Statement on inclusion

Our study was conducted by a research team composed entirely of scientists from the country where the study took place, all of whom share similar demographic characteristic. We acknowledge that this lack of diversity may have limited the range of perspectives represented in our work. We plan to address this limitation in future research by fostering more inclusive and diverse collaborations.

## Acknowledgements

MG, LFR and LS were supported by the PRIN Urbis grant (B53D23012400001) funded by the Italian Ministry for Universities and Research (MUR).

