## Supplementary Materials for "SANE: an Index of Anthropogenic Noise Levels for Wildlife Research in Terrestrial Ecosystems"

*Figure S1.* Map showing the distribution of the sampling points of this study. Land use data have been derived from Russo *et al*. (2025).

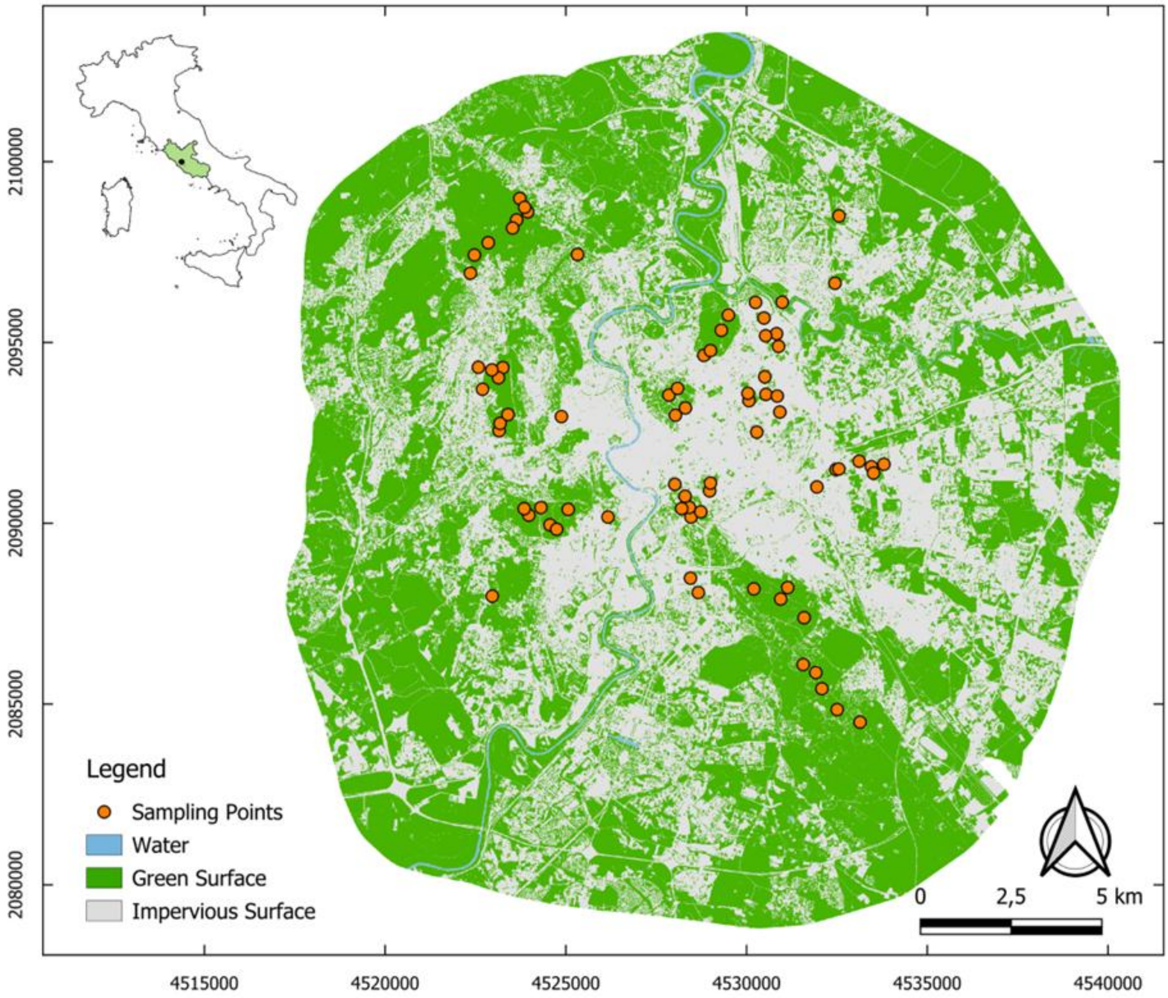

*Table S1*: Complete list of snippets used for simulated soundscapes creation. It includes the class, the specie and the source of each snippet.

| Class | Specie | Source |
|---|---|---|
| Human Noise | Human Voice | Colleagues in studio recordings |
| Human Noise | Siren | Sampled Recordings |
| Human Noise | High-revving Engine | Sampled Recordings |
| Human Noise | Power Tool | Sampled Recordings |
| Human Noise | Street Background Engine | Sampled Recordings |
| Human Noise | Brushcutter | Sampled Recordings |
| Anthrophonic-Birds | Common Pheasant | Elias A. Ryberg, XC364387. Accessible at www.xeno-canto.org/364387. |
| Anthrophonic-Birds | Common Raven | Simon Elliott, XC763188. Accessible at www.xeno-canto.org/763188. |
| Anthrophonic-Birds | Eurasian Curlew | Lars Edenius, XC803407. Accessible at www.xeno-canto.org/803407. |
| Anthrophonic-Birds | Eurasian Jay | Olivier SWIFT, XC858101. Accessible at www.xeno-canto.org/858101. |
| Anthrophonic-Birds | Hooded Crow | Johannes Worona, XC1006453. Accessible at www.xeno-canto.org/1006453. |
| Anthrophonic-Birds | Yellow-legged Gull | Théo Trees, XC899528. Accessible at www.xeno-canto.org/899528. |
| Anthrophonic-Birds | Magpie | David Bissett, XC999018. Accessible at www.xeno-canto.org/999018. |
| Anthrophonic-Birds | Grey Heron | Paul Kelly, XC936092. Accessible at www.xeno-canto.org/936092. |
| Non-Anthrophonic Birds | Italian Sparrow | Mats Rellmar, XC907069. Accessible at www.xeno-canto.org/907069. |
| Non-Anthrophonic Birds | Short-toed Treecreeper | João Tomás, XC944710. Accessible at www.xeno-canto.org/944710. |
| Non-Anthrophonic Birds | Crested Lark | Maarten Sluijter, XC950753. Accessible at www.xeno-canto.org/950753. |
| Non-Anthrophonic Birds | Long-tailed Tit | Jarek Matusiak, XC951570. Accessible at www.xeno-canto.org/951570. |
| Non-Anthrophonic Birds | Blue Tit | Jarek Matusiak, XC964477. Accessible at www.xeno-canto.org/964477. |
| Non-Anthrophonic Birds | Common Firecrest | Xavier Riera, XC986328. Accessible at www.xeno-canto.org/986328. |
| Non-Anthrophonic Birds | Common Redstart | Abe Bakker, XC994936. Accessible at www.xeno-canto.org/994936. |
| Non-Anthrophonic Birds | Eurasian Robin | Gonzalo Peña Sánchez, XC998115. Accessible at www.xeno-canto.org/998115. |

| | | |
|---|---|---|
| Non-Anthrophonic Birds | Eurasian Wren | Olivier SWIFT, XC1011343. Accessible at www.xeno-canto.org/1011343. |
| Non-Anthrophonic Birds | Eurasian Blackcap | Elias A. Ryberg, XC1017001. Accessible at www.xeno-canto.org/1017001. |

*Figure S2*: Variability of acoustic indices across simulated soundscapes. The 10th–90th percentile ranges are illustrated. The extension of the ribbon indicates the consistency of indices through the 100 simulated soundscapes.

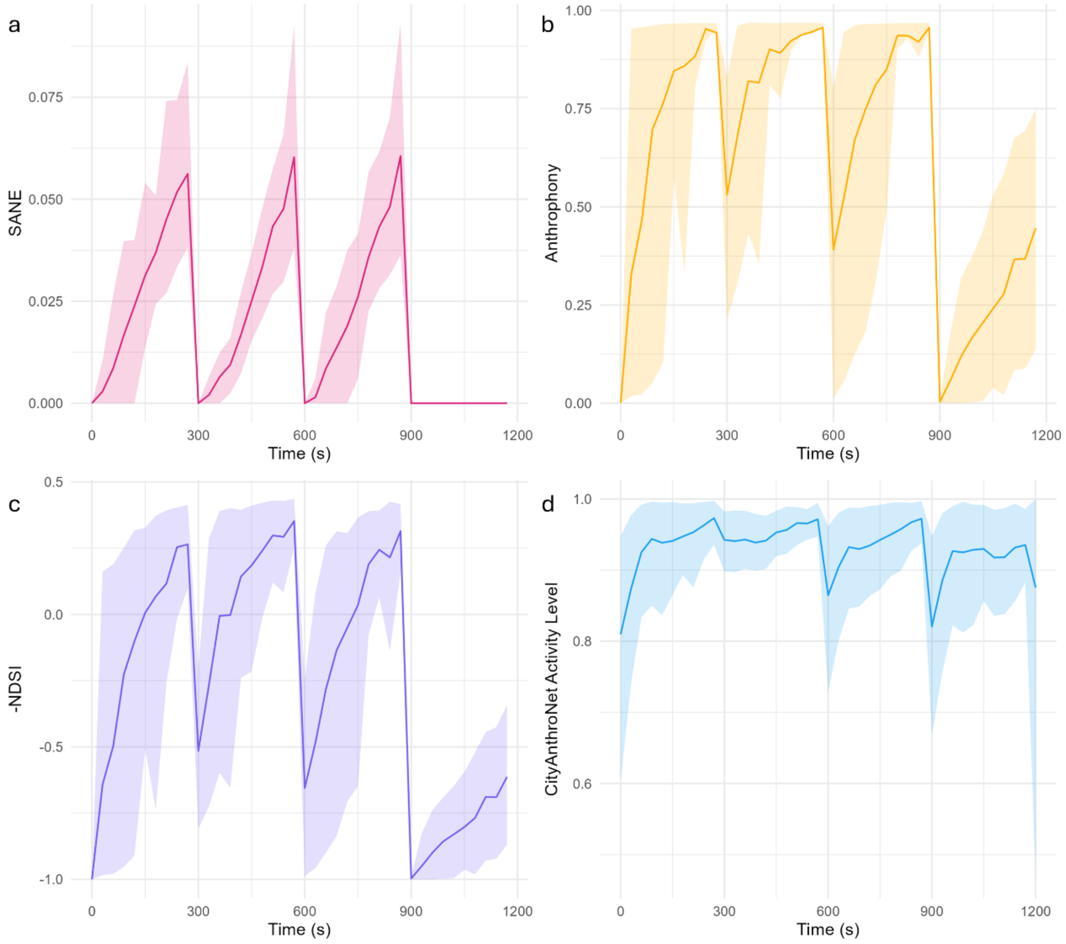
